# Improving Local Ancestry Inference through Neural Networks

**DOI:** 10.64898/2026.03.11.711082

**Authors:** Jazeps Medina Tretmanis, María C. Ávila-Arcos, Flora Jay, Emilia Huerta-Sanchez

## Abstract

**Motivation:** Local Ancestry Inference (LAI) allows us to study evolutionary processes in admixed populations[1], uncover ancestry-specific disease risk factors[2], and to better understand the demographic history of these populations[3]. Many methods for LAI exist, however, these methods usually focus on cases of intercontinental admixture. In this work, we evaluate both existing and novel methods in challenging scenarios, such as downsampled reference panels, intracontinental admixture, and distant admixture events.

**Results:** We present four novel LAI implementations based on neural network architectures, including Bidirectional Long Short-Term Memory and Transformer networks which have not previously been used for LAI. We compare these novel implementations to existing methods for LAI across a variety of scenarios using the 1 Thousand Genomes dataset and other synthetic datasets. We find that while all networks achieve high performance for intercontinental admixture scenarios, inference power is comparatively low for scenarios of intracontinental or distant admixture. We further show how our implementations achieve the best performance of all methods through specialized preprocessing and inference smoothing steps.

**Availability:** All implementations and benchmarking code available at https://github.com/Jazpy/LAINNs.

## 1 Introduction

Local Ancestry Inference (LAI) is a method used to assign population labels to segments of an individual’s genome by comparing sequence similarities between the unlabeled genome and a reference set of labeled genomes[4]. The goal of LAI is to facilitate the study of evolutionary processes in admixed populations[1], uncover ancestry-specific disease risk factors[2], and to better understand the demographic history of these populations[3]. In the Americas, European colonization and the transatlantic slave trade brought individuals of European and African descent who subsequently mixed with the Indigenous peoples. Today, many individuals from the Americas carry a combination of Indigenous, European, and African ancestries[5]. To understand the impacts of these recent admixture events, we need to infer local ancestries along the genome. By leveraging a reference panel consisting of genomes from Native American, African, and European populations, one can use LAI to partition the segments of an admixed individual’s genome by determining the reference population with the highest affinity for each segment and then labeling the segment accordingly. The availability of putatively unadmixed genome datasets[6] makes this task especially well suited for supervised learning approaches, and therefore all of the recent advances in machine learning methods can also be applied to LAI.

One prominent approach for LAI methods[7][8][9] is to build on Li and Stephens’ model[10], often implemented as Hidden Markov Models (HMMs), which label SNPs based on the most likely reconstruction from a set of observed labeled SNPs; an example of this approach is Recomb-Mix[11], which models LAI as a graph optimization problem. Alternatively, many other classification approaches can be used for LAI. For many years, RFMix[12] has been regarded as the gold standard for benchmarking LAI methods. RFMix first splits genomes into non overlapping windows (usually 1,000 SNPs) and then trains random forests independently on the raw SNP data of each genomic window. This “model-per-window” approach is effective because there is minimal correlation between the ancestry of distant genomic segments. Recent methods like G-Nomix[13] and LAI-Net[14] have also adopted this model-per-window approach, but have used other machine learning algorithms instead of random forests, while maintaining the same overall approach. Methods such as G-Nomix[13] and LAI-Net[14] have demonstrated improvements over RFMix in both accuracy and training time. SALAI-Net[15] provides an alternative by training on genetic distances between unlabeled and reference sequences, instead of just the raw SNPs. This means that, by encoding information from the reference and query sequences in the input, one model can be trained on genomic windows from different regions or even across species.

These methods opt for machine learning models different to the random forests used by RFMix, such as boosted trees, SVMs, and neural networks. Thus far, multilayer perceptrons (MLPs, used by LAI-Net)[14] and Convolutional Neural Networks (CNNs, used by SALAI-Net)[15] have been the primary neural network architectures applied to LAI tasks. These methods have been shown to have good LAI accuracy, but mostly in the specific scenario of recent intercontinental admixture. For example, it is unknown how accuracy is affected when i) the size of the reference panel is small, ii) the source populations are closely related, and iii) the admixture time is not too recent (older than 100 generations). These three points need to be considered because these are actual scenarios in real world applications. For example, it is common for unadmixed Native American reference populations to be composed of less than 100 individuals[16][17]. Another limitation is the lack of evaluating scenarios where reference populations belong to neighboring geographical regions with recent divergence times, such as Indigenous populations in America[11], consequently creating a gap in understanding the performance of LAI methods for admixed populations derived from closely related source populations. This intracontinental scenario should not be overlooked. While LAI on intercontinental datasets provides information about human evolution in the more distant past applying the same methods to a finer geographical resolution can yield information about human evolution for more recent timescales[3]. Finally, most evaluations of current methods assume a relatively recent admixture event (i.e., fewer than 50 generations ago)[12][13][14]. This means that LAI methods are rarely tested on individuals whose ancestry tracts have been fragmented into shorter, more dispersed segments over longer timeframes. Analyzing very short ancestry tracts, instead of potentially dismissing them as statisticla noise, could reveal previously unknown distant demographic events[18][19].

In this work, we evaluate the performance of current LAI methods[12][13][14][15][11] and four new LAI approaches based on fully connected, convolutional, recurrent, and transformer networks that we introduce here. We compare the existing LAI methods and our four new implementations under a variety of scenarios using both real data from the Thousand Genomes Project (1KG) dataset[6], as well as synthetic data simulated using msprime[20]. Specifically, we cover the following scenarios: (i) Intercontinental LAI using large reference panels from the 1KG (i. e., a simple, “baseline” scenario). (ii) Intercontinental LAI using downsampled reference panels from the 1KG.

(iii) Intracontinental (increasing sequence similarity between the source populations) LAI on recently admixed individuals, using European 1KG haplotypes as the base for our simulations. (iv) Intracontinental LAI on admixed individuals that represent admixed populations in America. (v) Intercontinental LAI on admixed individuals after simulating an old admixture event (reducing the length of the ancestry tracts). We show that LAI methods are very robust even when the samples sizes in the reference panel are smaller than 50 individuals. We find that current LAI methods underperform when the reference populations are genetically close, or when inferring ancestries with very short average tract lengths. In cases where standard approaches underperform, we achieve significant improvement gains by adding pre- and post-processing steps to our new LAI implementations. In the case of intracontinental admixture, a preprocessing step that highlights private variation leads to our implementation achieving significantly higher accuracy over existing methods in scenario (iv). Similarly, when inferring the ancestry of very short tracts, we achieve considerably higher accuracy over existing methods by complementing our LAI methods with a CNN-based smoother that is aware of distant admixture events. Our new LAI approaches will be useful for inferring genetic ancestry along the genome from both recent and more ancient admixture events.

## 2 Methods

### 2.1 Benchmarked Methods

We benchmarked a combination of both existing and new LAI methods (Table 1). Previously published methods were chosen to showcase different paradigms for LAI (e.g., “model-per-window” approach by RFMix, LAI-Net, and G-Nomix, graph-based by Recomb-Mix, and CNNs by SALAI-Net). Similarly, we designed four new approaches to showcase different neural network architectures. Specifically, using MLPs, CNNs, BLSTMs (Bidirectional Long Short Term Memory), and Transformers.

**Table 1:** List and description of benchmarked methods. We include information about which models are based on a neural network architecture, as well as marking architectures that have been built specifically for this work with ⋆.

| Method Name | Paradigms | Previously Published |
| --- | --- | --- |
| RFMix[12] | Model-per-window | Yes |
| G-Nomix[13] | Model-per-window | Yes |
| LAI-Net[14] | Model-per-window (Neural Network) | Yes |
| Recomb-Mix[11] | Graph-based, Haplotype reconstruction | Yes |
| SALAI-Net[15] | Position-agnostic sequence similarity (Neural Network) | Yes |
| MLP | Model-per-window (Neural Network) | No ★ |
| CNN | Model-per-window (Neural Network) | No ★ |
| BLSTM | Model-per-window (Neural Network) | No ★ |
| Transformer | Model-per-window (Neural Network) | No ★ |

We developed LAI models using four different neural network architectures. The first two architectures were used as sanity checks to compare against existing methods: MLPs, given their use in LAI-Net[14] and CNNs, for their previous use in SALAI-Net[15]. The last two architectures we included are BLSTM and Transformer architectures. These architectures were chosen for their specific use in sequential data, which maps well to analyzing SNP sequences. These architectures are commonly used in natural language processing tasks, which operate on data that share some concepts with SNP sequences. In both cases, the relative position of words or SNPs and local correlations within a window play a key role in classification.

Using these neural network architectures, we built a new “model-per-window” LAI method for each distinct architecture (Table 1). Details for these implementations are described in Section 2.6 “New LAI Architectures”. Before outputting the final ancestry predictions for these new implementations, we include a post-processing step on the raw outputs (Described in detail in Section 2.4). This is commonly known as a “smoothing” step, and its purpose is to remove statistical artifacts by leveraging predictions for surrounding regions before creating the final output since ancestries of adjacent regions are highly correlated.

Default settings were used for all previously published methods. For new methods (Table 1), we used a simple hyperparameter search optimization when training on dataset A (Table 2). For our hyperparameter search, we tried all combinations of candidates for two hyperparameters: batch size and learning rate, that were respectively chosen from the sets {16, 32, 64, 128} and {5×10^−3^, 1×10^−3^, 5×10^−4^, 1×10^−4^, 5×10^−5^, 1×10^−5^, 5×10^−5^, 1×10^−5^}. These hyperparameters control how many examples are given to the model and how much the model parameters can change at each training step. The selected hyperparameters were then used for training on all datasets. All new methods are based on a “model-per-window” approach, where the length of each window is always 1024 SNPs, which is a common default size used by other “model-per-window” approaches[12][14].

**Table 2:**
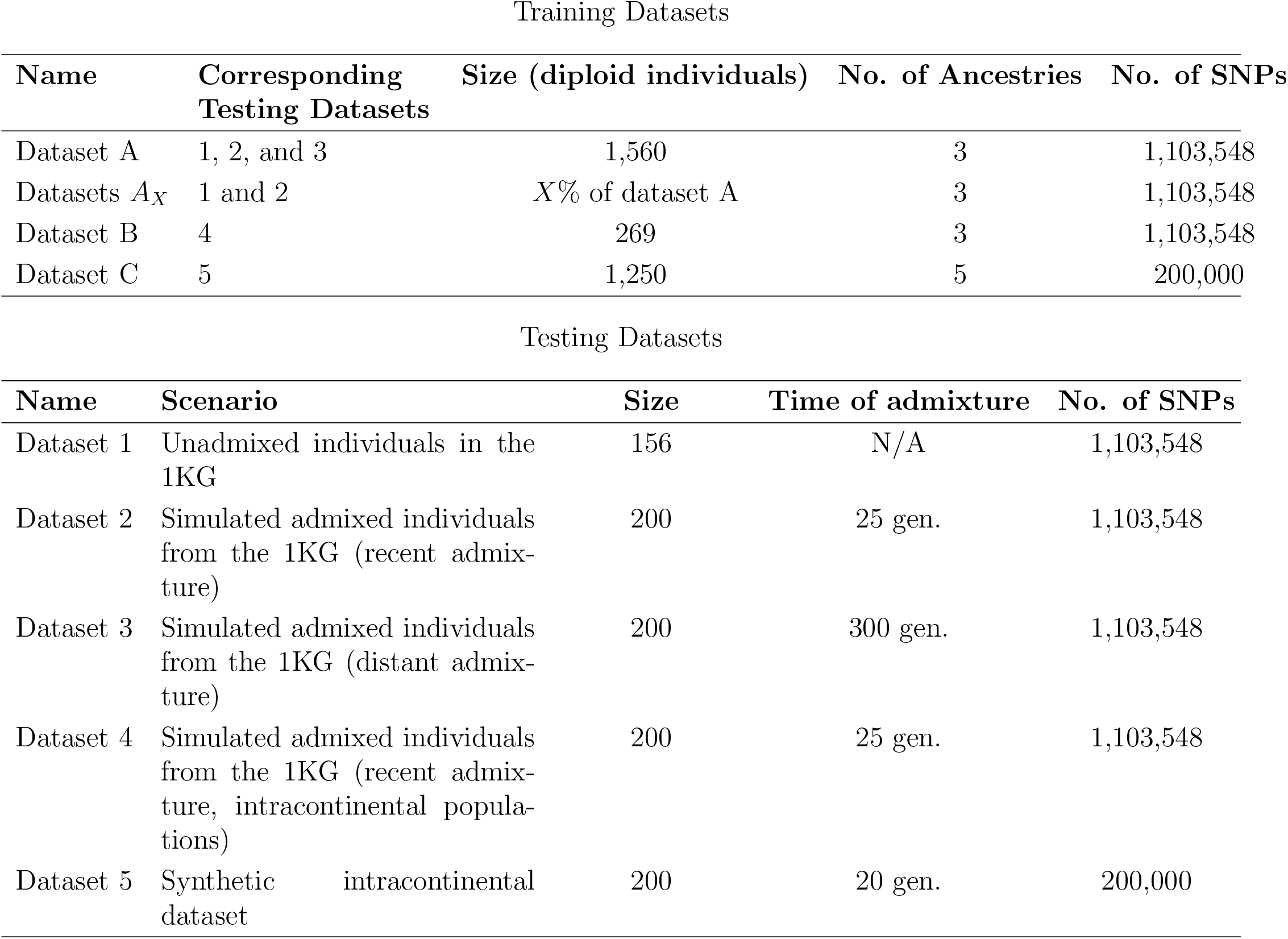
Scenario descriptions and sample sizes for all training and testing datasets. Datasets A, B, 1, 2, 3, and 4 are all created using real genomes from the 1KG. Datasets C and 5 which correspond to the 5-population intracontinental scenario are created entirely from synthetic genomes through *msprime*

Training Datasets
| Name | Corresponding Testing Datasets | Size (diploid individuals) | No. of Ancestries | No. of SNPs |
| --- | --- | --- | --- | --- |
| Dataset A | 1, 2, and 3 | 1,560 | 3 | 1,103,548 |
| Datasets $A_X$ | 1 and 2 | $X\%$ of dataset A | 3 | 1,103,548 |
| Dataset B | 4 | 269 | 3 | 1,103,548 |
| Dataset C | 5 | 1,250 | 5 | 200,000 |

Table 2: Scenario descriptions and sample sizes for all training and testing datasets.
| Name | Scenario | Size | Time of admixture | No. of SNPs |
| --- | --- | --- | --- | --- |
| Dataset 1 | Unadmixed individuals in the 1KG | 156 | N/A | 1,103,548 |
| Dataset 2 | Simulated admixed individuals from the 1KG (recent admixture) | 200 | 25 gen. | 1,103,548 |
| Dataset 3 | Simulated admixed individuals from the 1KG (distant admixture) | 200 | 300 gen. | 1,103,548 |
| Dataset 4 | Simulated admixed individuals from the 1KG (recent admixture, intracontinental populations) | 200 | 25 gen. | 1,103,548 |
| Dataset 5 | Synthetic intracontinental dataset | 200 | 20 gen. | 200,000 |

### 2.2 Training and testing datasets

#### Real genome-based intercontinental datasets

Training dataset A along with testing datasets 1, 2, and 3 (Table 2) correspond to intercontinental admixture scenarios. Training dataset A and testing datasets 1, 2, and 3 were constructed from the Thousand Genomes Project (1KG) dataset[6] using chromosome 22. We extracted 1, 560 unadmixed individuals in total with African (AFR; *n* = 553), European (EUR; *n* = 503), and East Asian (EAS; *n* = 504) continental ancestries. From these 1, 560 individuals, we randomly sectioned them into two sets of 1, 404 and 156 individuals. The set of 1, 404 (AFR *n* = 505, EUR *n* = 458, EAS *n* = 441) unadmixed individuals is taken as-is for training dataset A. The set of 156 (AFR *n* = 48, EUR *n* = 45, EAS *n* = 63) unadmixed individuals is taken as-is for testing dataset 1 (unadmixed individuals). For testing datasets 2 and 3 (admixed individuals), we perform a forward-in-time admixture simulation between these 156 unadmixed individuals using *haptools*[21] and a human recombination map of Chromosome 22[22]. Testing dataset 2 corresponds to 25 generations of admixture with initial proportions 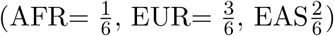. Testing dataset 3 corresponds to 300 generations of admixture with initial proportions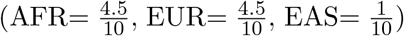. These proportions were chosen so as to have one underrepresented population (EAS), resulting in fewer tracts of this ancestry surrounded by the other 2 ancestries, making the smoothing process harder (see Section 2.4).

#### Real genome-based downsampled datasets

We also explore the effects of smaller reference panel sizes on LAI performance. This is representative of real-world scenarios where reference panels of understudied populations are common[16][1][17]. Therefore, it is of interest to measure how model accuracy is negatively impacted by decreasing panel sizes. For this we build eight datasets *A*_10_, *A*_20_, *A*_30_, …, *A*_80_, which are randomly downsampled versions of dataset A corresponding to 10%, 20%, 30%, …, 80% of the full dataset. The performance of models trained on these datasets is then evaluated by testing on datasets 1 and 2 (unadmixed and recently admixed individuals).

#### Real genome-based intracontinental datasets

Training dataset B and testing dataset 4 (Table 2) correspond to intracontinental tests performed on data extracted from the 1KG dataset. These are constructed identically to training dataset A (unadmixed individuals in the 1KG) and testing dataset 2 (forward-in-time simulation of admixture with individuals from the 1KG). However, instead of considering all 1, 560 individuals from Africa, Europe, and East Asia in the 1KG, we only consider European individuals of GBR (*n* = 91), FIN (*n* = 99), and TSI (*n* = 107) ancestries.

#### Synthetic intracontinental datasets

In order to test the methods on an even more difficult scenario, where our source populations are closely related and themselves admixed, we created training dataset C and testing dataset 5 (Table 2). We created purely synthetic genomes representing a Latin American demographic history using msprime[20] and the demographic model described by Medina-Muñoz et al.[23], This model presents a comprehensive demographic history for real-world populations in the 1KG[6] and unadmixed Indigenous Mexican populations[23]. We simulate 300 synthetic individuals which serve as stand-ins for 5 real-world populations: MXB (Unadmixed Native Americans in the Mexican Biobank[24]), MXL (Mexicans in the 1KG), CLM (Colombians in the 1KG), PEL (Peruvians in the 1KG), and PUR (Puerto Ricans in the 1KG). These 5 populations serve as the source populations for dataset C, compared to the 3 populations used for datasets A and B (Table 2). For each of these 5 synthetic populations, we randomly take 250 individuals to construct training dataset C. The remaining 50 individuals from each population are fed into *haptools*[21] for a forward-in-time admixture simulation of 20 generations with a uniform ^1^ initial proportion for each population.

### 2.3 Measuring accuracy

When measuring the performance of each method, it is important to specify that not all models output the same kind of information. Recomb-Mix and SALAI-Net output a per-SNP ancestry classification, while all other benchmarked methods (G-Nomix, LAI-Net, RFMix, and our new LAI implementations) output a per-window classification. When measuring accuracy, we count the number of correctly classified SNPs and divide by the total number of SNPs in the dataset. This means that for methods which output one per-window prediction, we treat all SNPs in that window as having the same ancestry prediction. We measure accuracy independently for each individual in all of the testing datasets, this allows us to also plot the standard deviation of the accuracy across the testing individuals whenever we plot results for any method.

When plotting accuracy results in Section 3, we do not differentiate between accuracies for specific ancestry tracts (i. e., plotting different accuracy measures for AFR or EUR tracts).

### 2.4 Smoothing

We implemented two different smoothing algorithms for the novel methods described in this work.

The first smoothing method is a weighted mode smoother. Each novel method outputs *c* values for every window, where *c* represents the possible number of ancestries we can use to classify SNP windows. These values correspond to likelihoods (adding up to 1) that the model assigns to each ancestry. To calculate the final prediction for a window with index *w*, we examine the surrounding windows and compute a weighted mode of the ancestries, using the raw probabilities as weights. This weighted mode is derived from the non-overlapping windows within the range of indices [(w - 12), (w + 12)]. We tested multiple lengths of the surrounding context for this smoothing method, using training dataset A and testing dataset 2 (simulated admixed individuals from the 1KG, see Table 2), with a radius ranging from 5 windows, to 20 windows. Using a radius of 12 yielded the best results on testing dataset 1 according to the accuracy metric described in Section 2.3. This smoothing method is applied to testing datasets 1, 2, 4, and 5.

Let *p*_*wa*_ denote the likelihood assigned to ancestry *a ∈* {1, …, *c*} for window *w*. The final, smoothed ancestry *â*_*w*_ for window *w* is calculated as the weighted mode over a window of radius *r*:

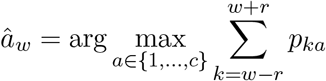

Where *r* is the smoothing radius (e.g., *r* = 12) and the sum aggregates the probabilities of each ancestry across the surrounding windows. The final prediction for window *w* is the ancestry with the largest total weighted probability within the range [*w* − *r, w* + *r*].

The second smoothing method is a neural network based on a U-Net architecture[25]. This smoother is exclusively used for testing dataset 3, as this smoother is specifically used to improve inferences on distant admixture datasets. This smoother takes as input the likelihoods produced by the model for all windows, represented in a (|*c*| + 1) × |*w*| matrix, where |*c*| is the number of ancestries and |*w*| is the number of windows (Figure 1). The first |*c*| rows of this matrix reflect the per-window probabilities for each ancestry, while the last row provides the genetic distance between a window and its predecessor (a value of 0 is used for the first entry in this row, as there is no predecessor). This information serves as a proxy for the probability of recombination between windows, which correlates directly with ancestry switches. We train this smoother using the true ancestries and model outputs from simulated datasets. These smoother training datasets are used solely for this purpose and are generated similarly to testing dataset 2 (Table 2), i. e., with the same original 1KG individuals, but with uniform initial ancestry proportions and admixture times set to 50, 100, 200, and 400 generations. Since these are independent simulations, ancestry switches occur at different positions, the ancestry tract length distributions vary, and the specific admixture time of 300 generations used in testing dataset 3 is excluded from the training data.

**Figure 1.**
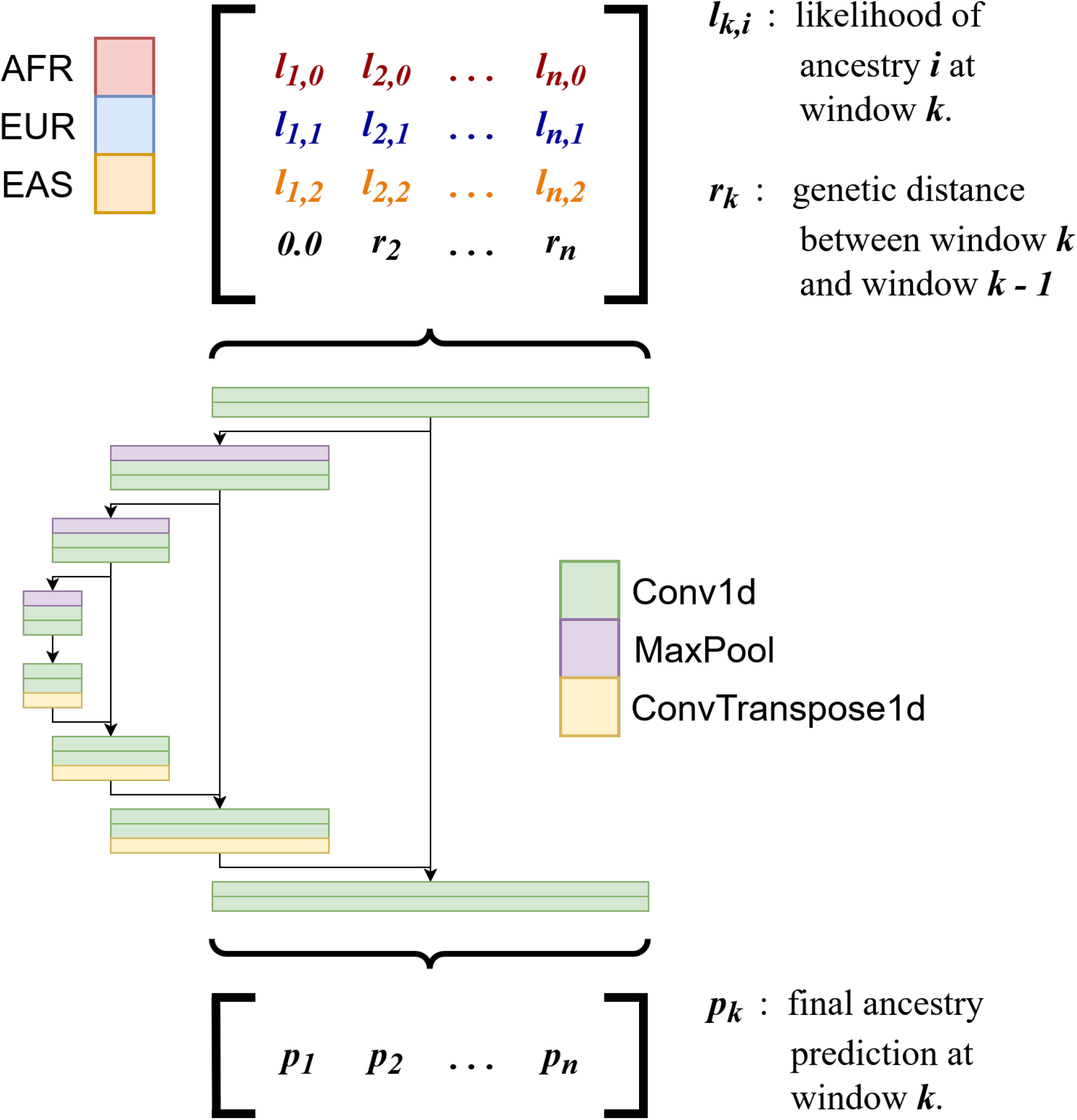
Diagramatic view of the U-Net Smoother. The input matrix is structured as follows: the first *c* rows correspond to the likelihoods of the possible source ancestries that can be inferred by the LAI model. The final row corresponds to the genetic distances between windows. Each column in this matrix represents a different, non-overlapping genomic window. This matrix is fed into a U-Net architecture with 4 downsampling and 4 upsampling layers. The final output is a one-dimensional vector that contains the final predictions for all windows.

### 2.5 Explicit encoding of private SNP information

When testing the new neural network implementations (see Table 1) on the intracontinental testing datasets (Testing datasets 4 and 5, Table 2), we also test a preprocessing step consisting of analyzing the training dataset (Training dataset B and C, Table 2) and finding SNPs where the derived allele is private to a single population, filtering for a minimum DAF *>* 1%.

After identifying the private SNPs for each population, we use this information to tag all SNPs for all sequences in both the training and testing datasets. We refer to the SNP information (i. e., the 0 or 1 information representing reference or alternative alleles) for position *l* in sequence *s* as *snp*_*ls*_. Each SNP’s tag *u*_*ls*_ can take on a value *u*_*ls*_ *∈* −1, 0, … *n*, based on the given SNP’s frequency across all *n* populations:

- *u*_*ls*_ = 0 if *snp*_*ls*_ is the ancestral allele,
- *u*_*ls*_ = −1 when the derived allele at position *l* is shared by multiple populations, and *u*_*ls*_ is the derived allele,
- *u*_*ls*_ = *n* when the derived allele at position *l* is private to population *n*, and *snp*_*ls*_ is the derived allele.

We derive *u*_*ls*_ for all SNPs (i. e., all positions for all individuals) by looking at both the individual’s alleles as well as the derived allele frequencies in all populations. After calculating all *u*_*ls*_ values, we use these instead of the original SNP sequence when feeding data into the network. Since we are calculating one tag for each SNP, we do not change the dimension of the data, just the representation of 0 or 1 for ancestral or derived allele. In principle, this approach is similar to other works that do not use raw SNP data, but information of private and shared SNPs across populations for the task of LAI[4][26]. A possible intuitive explanation for why this preprocessing step may be helpful is that providing private SNP information as an input feature makes the target function of the models easier to learn.

### 2.6 New LAI deep learning architectures

All of the new LAI architectures implemented in this work (Table 1) are based on the “model-per-window” paradigm. Under this paradigm, the genome is split into non-overlapping windows (in our case, we choose a length for these windows of 1, 024 SNPs). One model is then created for each of these genomic regions, and all models are independently trained on labeled windows for their associated region independently of each other. When labeling a query haplotype, we first section the haplotype, feed it independently into all of the trained models, and then “stitch” all of the results together to obtain the full labeling of the query haplotype. All of the methods we implemented for this work follow this procedure, with the only difference being the underlying neural network architectures used for each model. In the following subsections, we describe the architecture for each novel neural network in detail. All models presented in this study use the ReLU (Rectified Linear Unit) activation function.

#### Multilayer Perceptron (MLP)

This model is implemented with one input layer, two fully-connected layers, and one output layer. The number of neurons in each layer are assigned as follows: one neuron per SNP in the input layer (1, 024), twice the amount of neurons in the hidden layers (2, 048), and one neuron per possible source population in the output layer. The input and hidden layers are all followed by dropout layers with a proportion of 10%.

#### Convolutional Neural Network (CNN)

We implemented a one-dimensional convolutional neural network with four convolutional + average pooling blocks. The convolutional portion of these blocks is made up of two convolutional layers followed by batch normalization and a dropout layer with a proportion of 10%. Each pair of convolutional layers is followed by an average pooling step. The first of these blocks expects a one-dimensional input with only one channel of information, which corresponds to the input SNP sequence; the first convolutional block increases the number of channels to 8. Subsequent blocks increase the number of channels to 16, 32, and 64 channels respectively. In all cases, the convolution and pooling operations use a kernel size of 3.

Following these four convolutional + average pooling blocks, the output is flattened and passed through a fully connected classifier that has an analogous structure to the previously defined MLP.

#### Bidirectional Long Short-Term Memory (BLSTM)

The BLSTM architecture is implemented as three bi-directional LSTM layers stacked in sequence, the input length corresponds to the number of SNPs in a window (1, 024), and we define a hidden size for each of these BLSTM layers of 1, 024. Due to this bi-directionality, the output of the BLSTM layers is duplicated, resulting in an output tensor with 2, 048 values. This one-dimensional tensor is then fed into a fully connected classifier analogous to the MLP described above.

#### Transformer

For our Transformer-based architecture, we followed the standard encoder design proposed by Vaswani et al.[27]. The model processes 1D input SNP sequences of length 1, 024 and outputs source population likelihoods via a transformer encoder and a two-layer feedforward classifier.

We first obtain positional encodings for the input SNP sequence, these are sinusoidal positional encodings calculated from SNP positions within a specific window, not their physical position in the genome. The encoded input is passed through a 4-layer Transformer encoder, each with 4 heads of self-attention and feedforward sublayers. The output of the final encoder layer is passed through a feedforward decoder consisting of two linear layers: the first projects the transformer’s output dimensionality to half its original size, and the second maps it to the number of target classes.

## 3 Results

### 3.1 Unadmixed and intercontinentally admixed individuals based on the 1KG

We find that almost all models achieve a high accuracy (Figure 2), above 95%, for testing datasets 1 and 2 (Unadmixed and admixed intercontinental individuals in the 1KG, Table 2). This is expected, as this configuration of the data is a standard for previous LAI work [12][13][14][28][15][11]. Generally, performance for the unadmixed individuals is better than for the admixed individuals, this is also expected as most smoothing strategies will be helped by having only one contiguous ancestry tract, even if this is unrealistic. We find that G-Nomix and LAI-Net are the only methods to achieve higher accuracies in the admixed dataset than the unadmixed dataset. This could be attributed to their smoothing strategy, which explicitly accounts for admixed genomes[14]. We find that SALAI-Net achieves a noticeably lower accuracy than average (*∼* 93%). It is important to highlight that the key advantage of SALAI-Net is to be more flexible regarding differences in the training and testing data. It can, for example, perform inferences on a species different to the one it was trained on. When looking at the novel methods developed for this work, the Transformer architecture achieves the lowest overall accuracy for this dataset. Our MLP implementation achieves very similar accuracies to LAI-Net, which is expected as the neural network architecture is practically identical in both methods.

**Figure 2.**
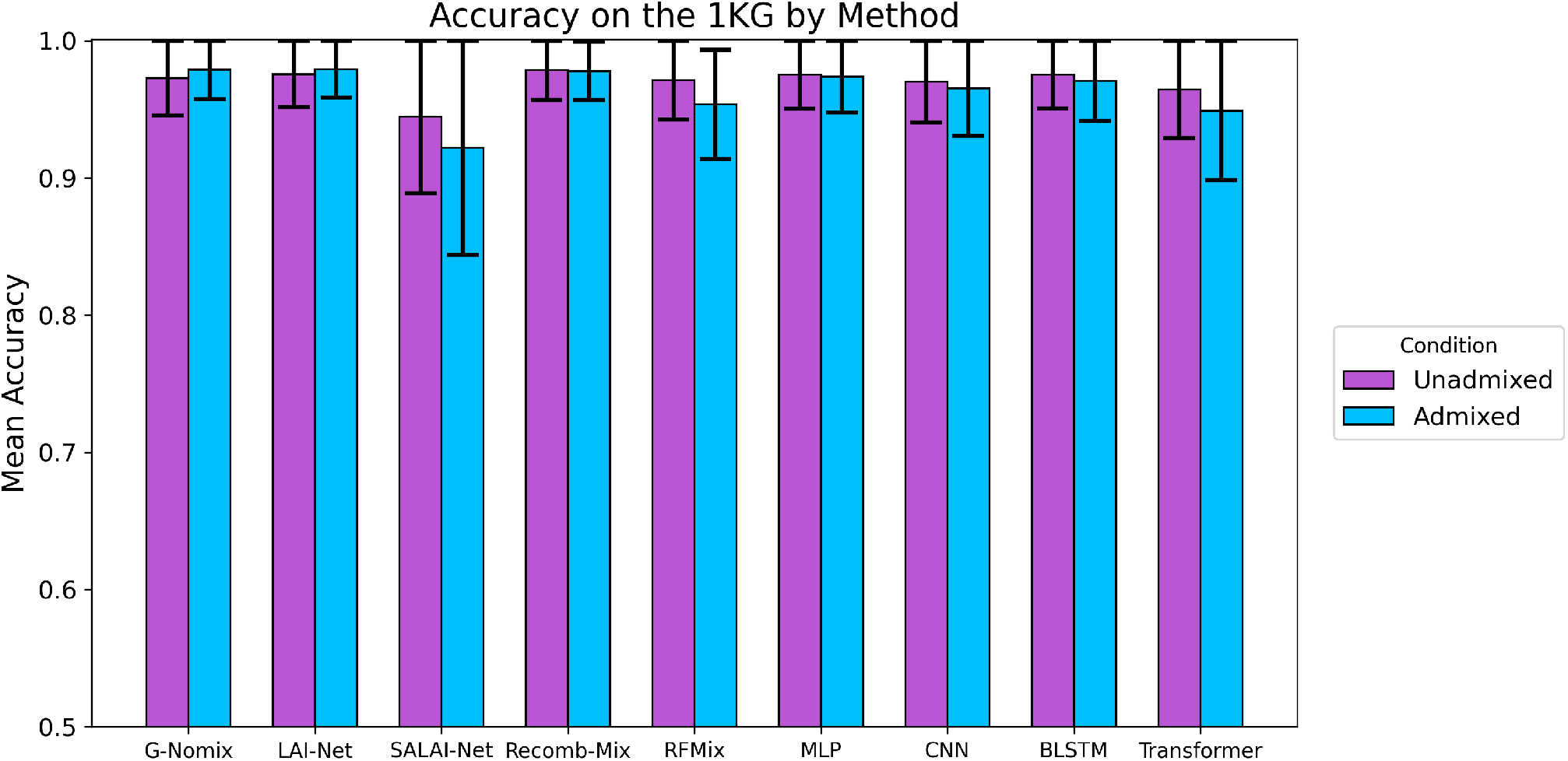
Admixed and unadmixed individuals in the 1KG (Datasets A, 1, and 2 in Table 2): LAI accuracy (y-axis) for the tested models (x-axis). Accuracies for testing dataset 1 (unadmixed) and dataset 2 (admixed through *haptools*) are shown in magenta and blue, respectively. Accuracies for the admixed testing dataset are generally lower than for the unadmixed dataset, especially for the Transformer architecture and SALAI-Net. In general, high accuracy is achieved by all methods.

### 3.2 Training on downsampled reference panels from the 1KG

In order to test the effects of smaller reference panels on accuracy, we repeated the methodology from Section 3.1, this time training on downsampled versions of training dataset A (Table 2). Figure 3 shows accuracies on unadmixed and admixed testing datasets 1 and 2 for all methods (Table 1). In general, we see very little loss in accuracy for the results on both the unadmixed and admixed testing datasets (Datasets 1 and 2, Table 2). This is the case even after downsampling to 10% of the original training dataset (Dataset A, Table 2), with less than 100 individuals per reference population.

**Figure 3.**
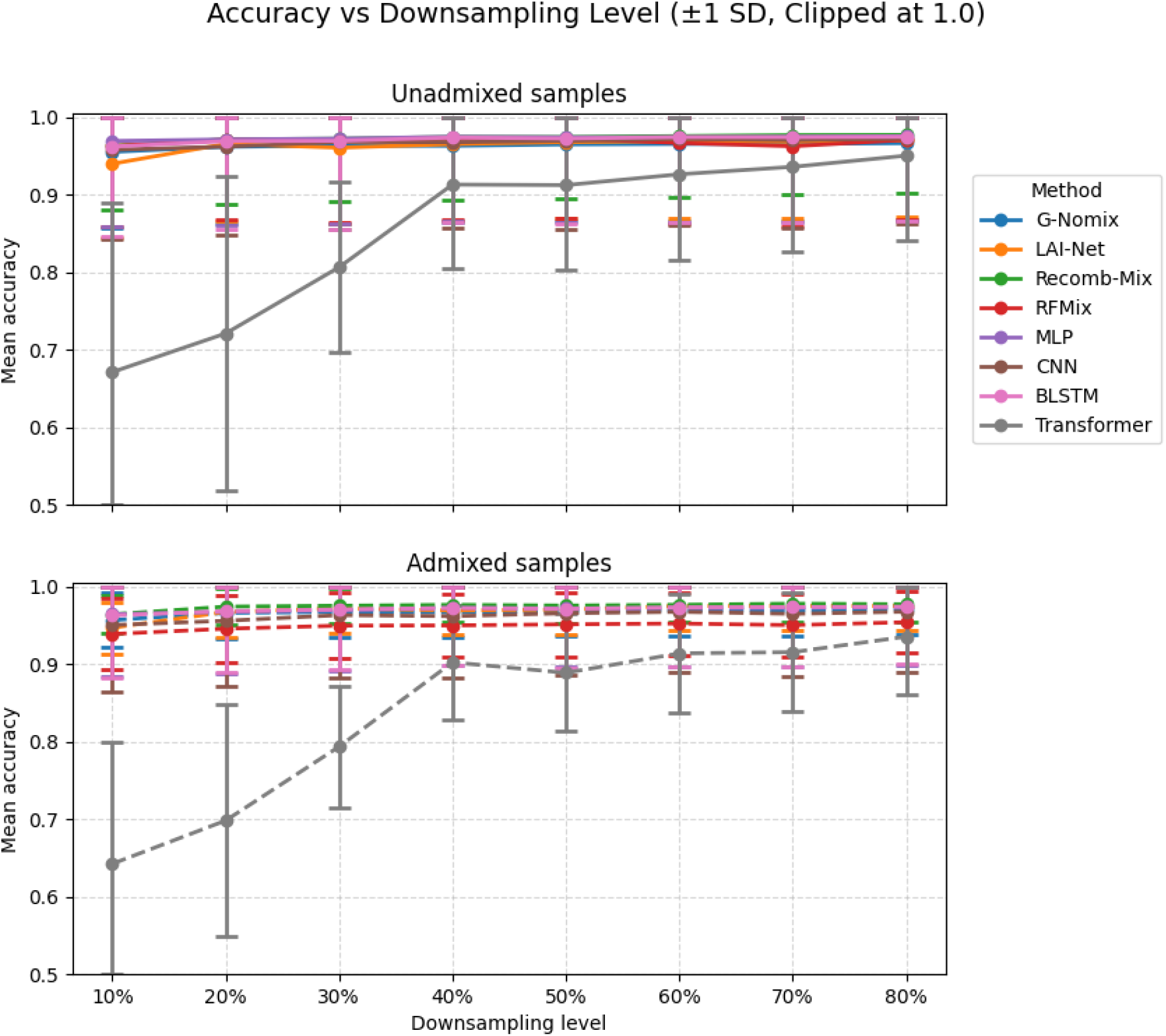
Admixed and unadmixed individuals in the 1KG (Datasets *A*_*X*_, 1, and 2 in Table 2): Training performed on downsampled versions of training dataset A (x-axis). LAI accuracy (y-axis) for the tested models (color). Upper plot shows results on the unadmixed testing dataset 1, and the lower plot shows results on the admixed testing dataset 2. In general, we see very little change in accuracy for all models, except for the Transformer architecture which sees a sharp drop-off in accuracy when reducing the training dataset to 30% of its original size or less.

The exception to this is the Transformer neural network architecture, which sees a sharp drop in accuracy and a large increase in the accuracy standard deviation as we reduce the size of of training dataset A to 30% or lower of its original size.

### 3.3 Intracontinentally admixed individuals based on the 1KG

After measuring model accuracies on the intercontinental scenario, we benchmarked the methods on populations that are more closely related (Figure 4). We created an intracontinental dataset using data from the 1KG, only considering individuals from GBR, FIN, and TSI ancestries which all belong to continental Europe (Table 2, Dataset B). Similarly to Section 3.1, we trained all models on unadmixed individuals belonging to these 3 populations, and tested on simulated admixed individuals in dataset 4 (Table 2). We find different trends to those in the intercontinental tests (Section 3.1). Recomb-Mix (70.25% accuracy) achieves the best results of the previously published methods benchmarked in this study, all methods show a marked decrease in accuracy compared to the intercontinental results. This is expected for multiple reasons. First, closely related populations will necessarily be harder to differentiate than populations that have drifted from each other for a longer period of time. Second, with the exception of Recomb-Mix, LAI methods are not commonly explicitly tested on intracontinental datasets. More importantly, we find that we can achieve a substantial increase in performance (59.35% vs. 72.32% accuracy) by preprocessing the data before feeding it into our CNN LAI implementation. This preprocessing step is described in Section 2.5, and shows that explicit encoding of private and shared alleles can be helpful for neural network-based LAI methods.

**Figure 4.**
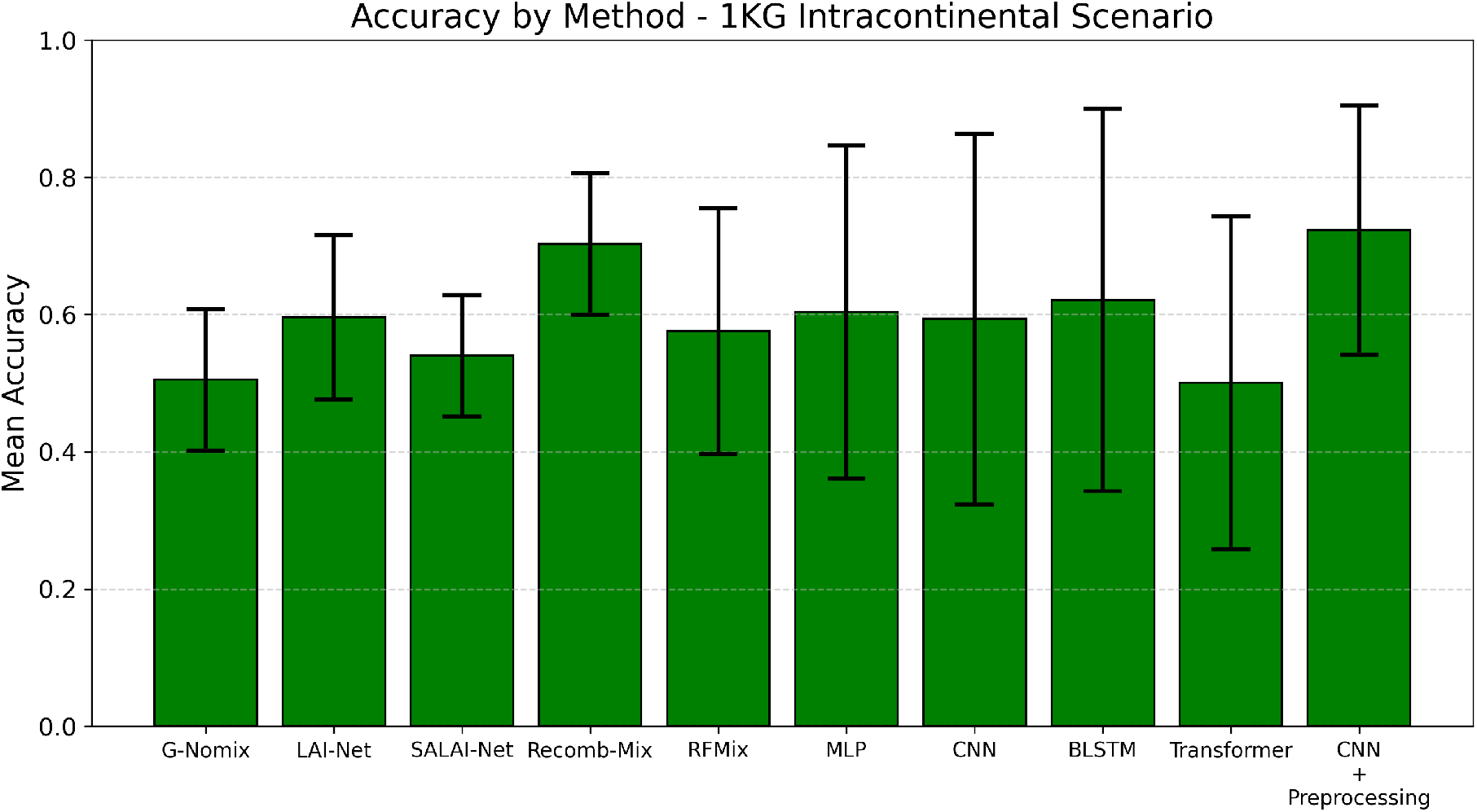
Intracontinental populations in the 1KG (Datasets B and 4 in Table 2): LAI accuracy (y-axis) for the tested models (x-axis). All models achieve markedly lower accuracies when compared to more standard datasets. Recomb-Mix achieves the highest accuracy of the published methods, while the highest overall accuracy is achieved by our CNN implementation after preprocessing the data to explicitly include private SNP information.

### 3.4 Intracontinental synthetic individuals

We tested an even more difficult intracontinental scenario (Figure 5), where the reference populations are stand-ins for real populations in continental America (Datasets C and 5, Table 2). This scenario is still of practical interest, as LAI inferences could be used to learn more about recent migrations in post-America. We created synthetic individuals representing 5 closely related populations in continental America to create training dataset C and testing dataset 5 (Table 2). We find that all models show a dramatically lower accuracy for this dataset (*<* 40% accuracy for all published models except Recomb-Mix and LAI-Net). This is similar to what we see for the other intracontinental dataset (Section 3.3), in general, all methods perform much worse, this is partly due to the increase in source populations (5 vs. 3), in addition to the added complexity of the source populations sharing many similarities due to their demographic history. Once again, we find that we can achieve a substantial increase in performance (34.60% vs. 62.77% accuracy) by preprocessing the data before inference by our CNN LAI implementation. When we include this preprocessing step, our CNN implementation achieves the highest accuracy of all models.

**Figure 5.**
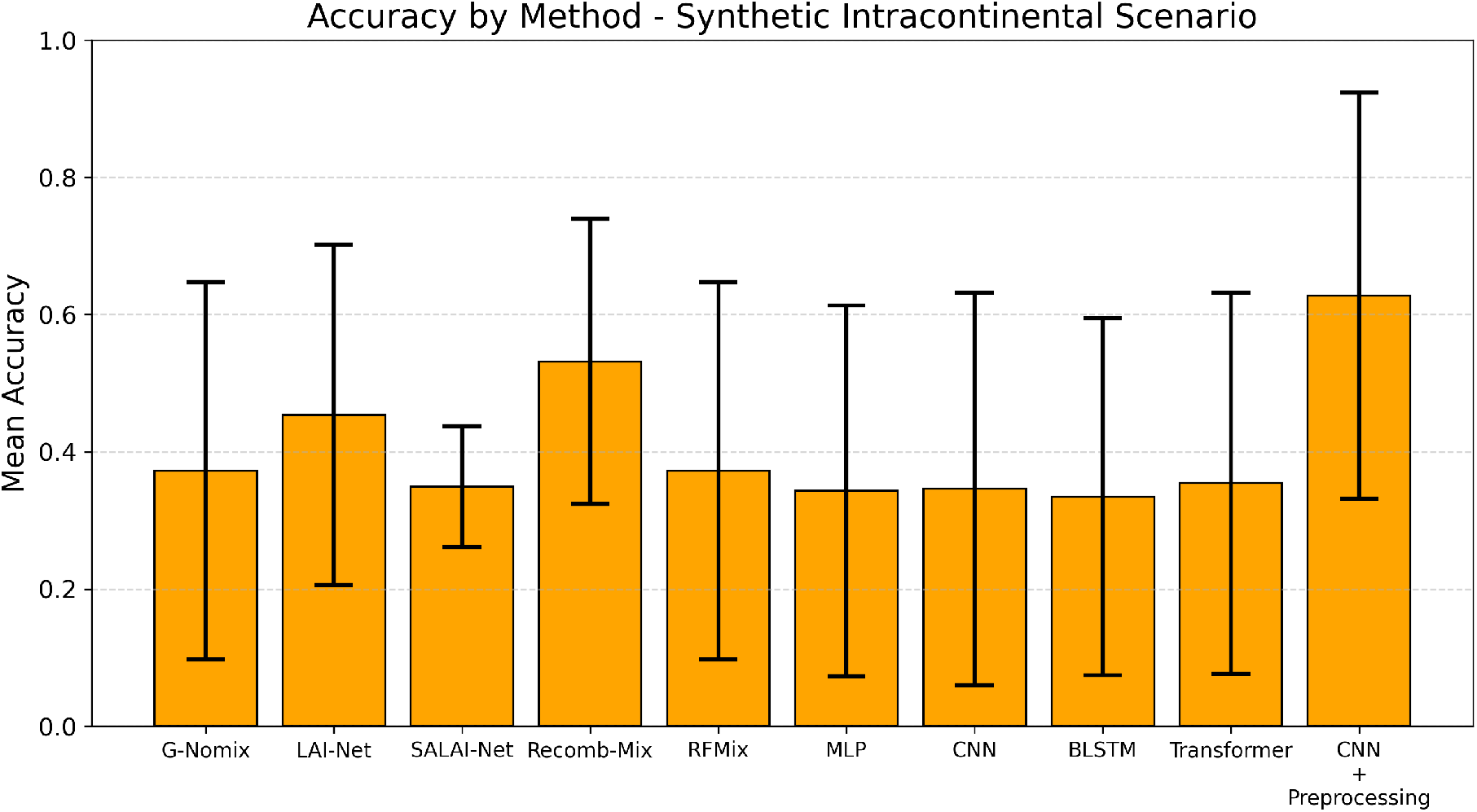
Synthetic intracontinental dataset (Datasets C and 5 in Table 2): LAI accuracy (y-axis) for the tested models (x-axis). All models achieve markedly lower accuracies when compared to more standard datasets. Recomb-Mix achieves the highest accuracy of the published methods, while the highest overall accuracy is achieved by the CNN implementation of LAI after preprocessing the data to explicitly include private SNP information.

### 3.5 Distant and intercontinental admixture

Finally, we benchmarked the existing methods along with our CNN implementation for LAI for histories involving older admixture events and with heterogeneous ancestry proportion (Figure 6) For these results, we plot the accuracy of each ancestry independently, as the recall of EAS tracts is of special interest due to its low proportion in the admixed population. We created a non-realistic scenario of admixture in the Americas, with 300 generations of admixture of 3 source populations: AFR, EUR, and EAS (Testing dataset 3, see Table 2). Furthermore, we set a low (10%) proportion for the EAS ancestry, resulting in short EAS tracts due to time of admixture and overall proportion. For this dataset (Testing dataset 3, Table 2), we used populations from the 1KG along with a forward-in-time simulator to create the artificially admixed individuals. Testing on this dataset shows that this is a challenging scenario for most methods, as some ancestry tracts will be sparsely distributed and very short when admixture happened deep in the past (*>* 100 generations). Figure 6 shows that accuracies on testing dataset 3 (Table 2) are strongly affected by what the true ancestry of a SNP is. All models are good at inferring the ancestry of AFR and EUR SNPs as these SNPs will belong to longer tracts that will be less likely to be discarded as statistical noise. Meanwhile, SNPs of EAS ancestry are inferred with a much lower accuracy, which is expected as EAS ancestry was simulated to appear with a proportion of only 10% throughout the genome. Simulating admixture for 300 generations further ensures that EAS ancestry tracts will have relatively short lengths on average. Of the published methods, we find that LAI-Net achieves the highest accuracy by a considerable margin. The highest accuracy for the underrepresented EAS ancestry is achieved by our baseline (no private SNP information) CNN implementation after coupling it with the neural network-based smoothing strategy described in Section 2.4. This is expected, because explicitly accounting for shorter tract distributions during smoother training makes the distant admixture dataset more similar to the data distribution expected by the model, decreasing the likelihood that short tracts are dismissed as statistical noise.

**Figure 6.**
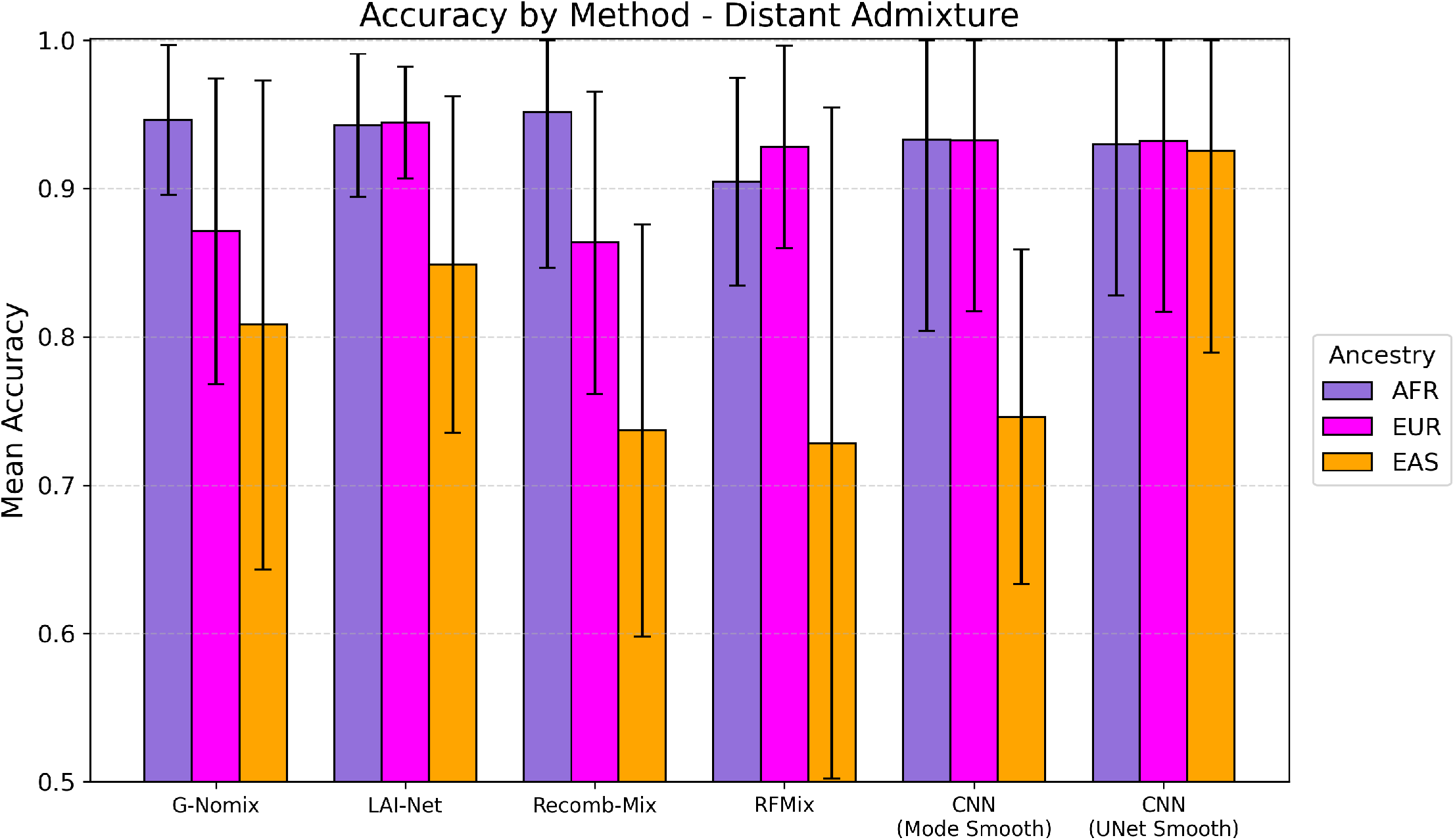
Distant admixture dataset (Datasets A and 3 in Table 2). LAI accuracy (y-axis) for the tested models (x-axis). *haptools* is used to simulate 1KG individuals after 300 generations of admixing between AFR, EUR, and EAS populations. Accuracies for AFR, EUR, and EAS tracts are shown in purple, magenta, and orange respectively. The highest accuracy for the underrepresented ancestry (EAS) is achieved by LAI-Net when considering only the published methods, the highest overall accuracy is achieved by our CNN implementation after implementing a distant admixture-aware smoother.

## 4 Discussion

In this study, we have benchmarked Local Ancestry Inference (LAI) using both existing methods and five new approaches introduced here. We find that, under the intercontinental scenario when admixture is recent and source populations are distant, LAI can be performed with very high accuracy (Figure 2) with all methods (Testing datasets 1 and 2, Table 2). This is especially true for methods that adopt the “model-per-window” approach as popularized by RFMix. Notably, this holds even when we consider cases when the size of the reference panels is as low as *∼* 50 individuals per population (Figure 3). However, when considering closely related intracontinental populations (Testing datasets 4 and 5, Table 2), or very short ancestry regions that originate from older (*>* 100 generations in the past) admixtures (Testing dataset 3, Table 2), we find that most of the previously published methods underperform. Specifically, our intracontinental datasets (Datasets B, C, 4, and 5 in Table 2) results, show that existing methods achieve markedly lower accuracies when populations are closely related (Figures 4 and 5). For the simulated intracontinental dataset with 5 populations (Datasets C and 5 in Table 2, Figure 5) this loss in accuracy is partly due to the increase in the number of source populations that make up the reference panel for the admixed individuals.

When testing on intracontinental datasets, the highest accuracy of previously published methods is achieved by Recomb-Mix for both datasets (49.34% and 70.25% accuracies). Possible reasons for this include the specific inclusion of intracontinental datasets in its original benchmarking[11], the way genetic distance is incorporated as a data input to account for recombination, or the fact that it uses a graph-based paradigm instead of the “model-per-window” framework for LAI. Importantly, the highest accuracy of all the benchmarked methods for both intracontinental datasets (5) is achieved by our CNN implementation after including a data preprocessing step (Section 2.5) that explicitly encodes which variants are private to only one population in the reference panel (62.77% and 72.32% accuracies). This preprocessing step guides the network by providing information of shared and private alleles that we know to be biologically relevant for LAI, and this is similar in principle to methods that leverage private variation to classify archaic hominid ancestry, such as HMMix[29] and DAIseg[26]. Recomb-Mix, the published method with the highest accuracy on the intracontinental datasets, also encodes some notion of private SNPs when constructing its haplotype graph, further showing that this is useful information for LAI.

We also consider the case of older admixture (*>* 100 generations) which is not commonly measured by the LAI methods studied here. This case is more challenging because we expect shorter tract-lengths which are more difficult to distinguish from statistical noise with typical smoothing strategies. We implemented a distant-admixture smoother and trained it with simulated data under a range of admixture times (25 to 400 generations), as well as including the genetic distance between ancestry windows as an input feature. This smoothing implementation achieves an accuracy of *∼* 92% which is higher than all the methods investigated in this study (*∼* 85% accuracy) (Figure 6). Applications of this implementation to human genomic datasets may reveal previously undetected admixture events, and support the hypothesis that admixture has been more common than previously thought.

While our LAI implementations perform best, there is still room for improving these approaches. In the case of intracontinental admixture, it may be that further preprocessing steps could increase the accuracy; future work could go beyond SNP sequence information and explore how inference is affected when models consider additional input features that capture the genetic relationships among the source populations. In general, we believe that neural networks are a powerful tool for the complicated feature space of ancestry deconvolution, and that further LAI performance gains may be achieved by the principled inclusion of data features derived from raw SNP data, instead of the use of larger, more computationally expensive models. While it is possible that much larger models could achieve similar improvements working on just SNP sequences, we value keeping model size and training time low. This is also reflected by our distant admixture scenario (Dataset 3, Table 2), where training the final ancestry smoother on data that more closely represents hard-to-infer short ancestry tracts improves performance.

Our results highlight the potential of neural network–based approaches for LAI, while also pointing to substantial opportunities for future work. For example, we do not consider archaic introgression in this study, although it is a closely related problem that can allow us to study how archaic introgression has impacted more recent admixture events[30], and could benefit from similar modeling and preprocessing strategies to the ones presented in this work.

## 5 Conclusions

In this work, we introduce novel neural network-based methods for Local Ancestry Inference and provide a comprehensive benchmark against existing approaches. To our knowledge, this is the first time that all these methods have been evaluated under identical conditions (same training and testing data sets and demographic scenarios). Our evaluation spans standard and challenging LAI scenarios, including downsampled reference panels, closely related intracontinental populations, and distant admixture events (*>* 100 generations). Not only are these results helpful for users when deciding which method best suits their use case, but we also show how the preprocessing of SNP data and the inclusion of recombination-aware smoothers result in considerable increases in accuracy for intracontinental and older admixture scenarios.

## 7 Data availability

The novel LAI implementations and benchmarking code are available on GitHub at https://github.com/Jazpy/LAINNs.

## 8 Acknowledgments

This work was supported by the Human Frontier Science Program, the National Institutes of Health, the Secretaría de Ciencia, Humanidades, Tecnología e Innovación, and the Agence Nationale de la Recherche.

